# Mitochondrial adenine base editing of mouse somatic tissues via adeno-associated viral delivery

**DOI:** 10.1101/2024.12.10.627690

**Authors:** Christian D. Mutti, Lindsey Van Haute, Lucia Luengo-Gutierrez, Isabel Dowling, Jose D. Barrera-Paez, Keira Turner, Pedro Silva-Pinheiro, Michal Minczuk

## Abstract

The development of adenine base editing in mitochondria, alongside cytidine base editing, has significantly expanded the genome engineering capabilities of the mitochondrial DNA. We tested the recent advancements in adenine base editing technology using optimised TALEs targeting genes *Mt-Cytb, Mt-CoII* and *Mt-Atp6* in mouse cells, and observed successful A:T to G:C conversions within the target windows of each gene. We then used the best-performing Mt-Atp6 pairs for systemic AAV9 delivery to neonatal mice and quantified editing in somatic tissues after 4 weeks and 6 months. Adenine editing was low at 4 weeks and increased only modestly after 6 months, indicating that prolonged exposure alone is insufficient to overcome the limited activity of the editor architectures tested here in vivo. These findings establish the feasibility of AAV-delivered mitochondrial A-to-G editing while defining important limitations that require further optimisation.

## Introduction

Mitochondria, essential organelles responsible for energy production and cellular homeostasis, possess their own genome distinct from the nuclear one. Mitochondrial DNA (mtDNA) alterations have been implicated in various human diseases, making precise genome editing tools invaluable for both basic research and potential therapeutic interventions [1]. Base editing has emerged as a transformative technology for precise genome manipulation, offering unprecedented capabilities for targeted nucleotide alterations without inducing double-strand breaks. While extensively explored in the nuclear genome, the application of base editing to mtDNA presents unique opportunities, but also challenges [2].

The inability to efficiently deliver nucleic acids into mitochondria has prompted the development of genome modification technologies based on custom DNA-binding protein. These technologies included mitochondrially-targeted zinc finger nucleases (mtZFNs), transcription activator-like effector nucleases (mitoTALEs) and meganuclease (mitoARCUS) and have been used for site-specific cleavage of mtDNA [3–11]. Combining ZF or TALE programmable DNA-binding proteins with deaminase enzymes, such as cytidine or adenine deaminases, allows for C:G to T:A or A:T to G:C conversions, respectively. The first major breakthrough was DddA-derived cytosine base editors (DdCBEs) which allowed for site-specific C:G to T:A edits in the mtDNA for the first time [12]. The initial dimeric DdCBE architecture contained the catalytic domain of DddAtox bacterial cytidine deaminase acting on double-stranded DNA, split into two inactive fragments (at positions G1333 or C1397), which were brought together by TALE DNA binding domains [12]. The technology was rapidly used by our lab and others for many applications including; knocking out each mtDNA-encoded protein-coding gene[13, 14], reverse genetics approaches [15] and *in vivo* applications in zebrafish [16, 17], rats [18] and mice [19, 20]. The technology also evolved to improve efficiency and widen compatibility for different sequence contexts and DNA-binding proteins [21–25]. The discovery of mitochondrial cytidine base editing provided foundation for the development of adenine base editors in mitochondria using the TadA8e enzyme, deemed transcription-activator-like effector (TALE)-linked deaminases (TALEDs), majorly increasing the capabilities of the mitochondrial gene editing toolkit [26]. Further developments included adenine editors containing a nickase to generate the single-stranded DNA required for deaminase activity (mitoBEs) [27]. In this study, we sought to apply the adenine base editing technologies, TALEDs and mitoBEs, *in vivo* using adeno-associated viral delivery to mouse somatic tissues.

## Materials and methods

### Plasmid construction and viral vectors

The TALE sequences and plasmids used were designed as reported in [13]. To construct the plasmids used in the cell screen, the functional domain was changed to varying architectures of TadA8e, DddAtox(E1347A) or BspD61(C) using AvrII and XbaI for pTracer or BamHI and KpnI for pcDNA3.1. Vector construction intended for AAV production was achieved by restriction digest of the transgenes using 5’ NotI and 3’ BamHI sites, allowing cloning into a rAAV2-CMV backbone, previously reported in [20]. The cloned plasmids were used to generate recombinant AAV9 viral particles at the UNC Gene Therapy Center, Vector Core Facility (Chapel Hill, NC) and VectorBuilder.

### Cell culture and transfections

NIH/3T3 cells (CRL-1658TM, obtained from the American Type Culture Collection (ATCC)) were maintained at 37 °C in a 5% (vol/vol) CO2 atmosphere in complete Dulbecco’s Modified Eagle Medium (DMEM) containing 4.5 g/L glucose, 2 mM glutamine, and 110 mg/ml sodium pyruvate, supplemented with 10% calf bovine serum with iron and 5% penicillin/streptomycin (all sourced from Gibco). Regular mycoplasma tests performed on the culture medium yielded negative results.

For the adenine base editor pair screening, NIH/3T3 mouse cells were seeded in six-well tissue culture plates at a 70% confluency and transfected with 3200 ng of each monomer (L and H), using 16 µl of FuGENE-HD (Promega) according to the manufacturer’s instructions. After 24 h, the cells were harvested for Fluorescence-activated cell sorting (FACS) and sorted for GFP and mCherry double-positive cells employing a BD FACSMelody^TM^ Cell sorter. The collected double-positive cells were allowed to recover for an additional 6 days before being utilised for DNA extraction, as detailed below. For cell survival experiments, AAV plasmids were transfected with empty GFP plasmid in a 1:1 ratio, sorted using BD FACSMelody^TM^ Cell sorter and replate for 3 days. Cells were then analysed using Countess™ 3 Automated Cell Counter (Invitrogen) to determine cell survival.

### Animals

(Ethics statement) All animal experiments were approved by the local Animal Welfare Ethical Review Body (AWERB) at the University of Cambridge and carried out in accordance with the UK Animals (Scientific Procedures) Act 1986 (Procedure Project Licence: PP1740969) and EU Directive 2010/63/EU). Mice on C57BL/6J background were sourced from Charles River Laboratories. Animals were housed in a temperature- and humidity-controlled environment, maintained under a 12-hour light/12-hour dark cycle, and continuous access to food and water *ad libitum*. Euthanasia was carried out via cervical dislocation. Newborn pups [both males and females on Postnatal day 1(P1)] were injected with a dose of 5 × 10^11^ vg per monomer per animal (1 x 10^12^ vg total) via the temporal vein (systemic administration), using a 30 G, 30° bevelled needle syringe. Control P1 pups were injected with equivalent volumes of vehicle buffer (1× PBS, 230 mM NaCl, and 5% w/v D-sorbitol). A total of 3 animals per group were used. The in vivo experiments were designed as molecular proof-of-concept studies, with mtDNA editing as the primary endpoint rather than as powered phenotypic comparisons. The relatively small group sizes therefore limit statistical power and the precision of estimates of tissue-specific editing efficiency. The 4-week cohort contained n = 3 animals per group; the 6-month cohort contained n = 4 control, n = 4 TALED and n = 3 mitoBE animals.

### Genomic DNA isolation and Sanger sequencing

NIH/3T3 mouse cells underwent trypsinisation for collection, followed by a single wash in PBS and extraction using the DNeasy Blood & Tissue Kit (QIAGEN) following manufacturers guidelines. The lysates were then extracted using DNeasy Blood & Tissue Kit (QIAGEN) following manufacturers guidelines. Genomic DNA extraction from mouse samples was also extracted using the DNeasy Blood & Tissue Kit (QIAGEN) following manufacturers guidelines. For Sanger sequencing, the edited region underwent PCR amplification using GoTaq G2 DNA polymerase (Promega) and using primers and PCR protocol for each site as listed in [13] PCR purification and subsequent Sanger sequencing were conducted by GENEWIZ/AZENTA (UK).

### High-throughput targeted amplicon mtDNA sequencing, processing and mapping

mtDNA-wide sequence analysis was performed, sequenced and analysed as in [13]. Briefly, two overlapping long amplicons (8,331 bp and 8,605 bp) spanning the entire mtDNA molecule were generated via long-range PCR utilising PrimeSTAR GXL DNA polymerase (TAKARA). Tagmentation and indexing PCR were performed using the Nextera XT index kit (Illumina, FC-131-1096) as per the manufacturer’s instructions. Subsequently, libraries were subjected to high-throughput sequencing on the Illumina NovaSeq platform (PE250), with demultiplexing performed using the respective manufacturer’s software.

For processing and mapping of the high-throughput data related to mtDNA-wide analysis, sequenced reads underwent quality trimming and 3′ end adaptor clipping simultaneously using Trim Galore! (--paired). Reads were aligned to ChrM of the mouse reference genome (GRCm39) with Bowtie2 (--very-sensitive; --no-mixed; --no-discordant). Count tables were then generated using samtools mpileup (-q 30) and varscan.

### Quantification of viral genomes copy number and relative mtDNA content by quantitative real-time PCR

Quantitative real-time PCR analyses were conducted in a QuantStudioTM 3 system (Thermo Fisher) using TaqMan™ Gene Expression Master Mix (Thermo Fisher), following the manufactureŕs instructions, and Ct values were obtained using the system’s built-in software. Each reaction was performed in a final volume of 20 µl in technical triplicates, with each dataset representing samples from an individual mouse.

To assess the relative mtDNA content in mouse tissue samples, mtDNA levels were quantified using primers and probes targeting MT-ND1: Mmu_Nd1_Fw: 5’-GAG CCT CAA ACT CCA AAT ACT CAC T −3’; Mmu_Nd1_Rv: 5’-GAA CTG ATA AAA GGA TAA TAG CTA TGG TTA CTT CA −3’; Mmu_Nd1_probe: 5’-/56-FAM/ CCG TAG CCC /ZEN/ AAA CAA T/3IABkFQ / −3’. MT-ND1 levels were normalized to the nuclear gene Actin using the following primers and probe: Mmu_Actin_Fw: 5’-CTG CTC TTT CCC AGA CGA GG −3’; Mmu_Actin_Rv: 5’-AAG GCC ACT TAT CAC CAG CC −3’; Mmu_Actin_probe: 5’-/56-TAMN/ ATT GCC TTT CTG ACT AGG TG /3BHQ/ −3’. ΔΔCt analysis was employed to calculate the relative mtDNA copy number, with vehicle-injected mice serving as the control.

### Immunoblotting of adenine base editors in mouse tissues

Mouse tissue samples (∼50 mg) underwent homogenization in 200 μL of ice-cold RIPA buffer (consisting of 150 mM NaCl, 1.0% NP-40, 0.5% sodium deoxycholate, 0.1% SDS, and 50 mM Tris, pH 8.0) supplemented with 1X cOmplete^TM^ mini EDTA-free Protease Inhibitor Cocktail (Roche, UK), utilizing a gentleMACS^TM^ dissociator. Following homogenization, the samples were incubated on ice for 20 mins and then clarified by centrifugation (20,000 × g for 20 min at 4 °C). Protein lysates (∼20 μg) were combined with 10 X NuPAGE™ sample reducing agent and 4X NuPAGE™ LDS sample buffer (Invitrogen), followed by incubation for 5 min at 95 °C. The denatured protein samples were subsequently separated in a Bolt 4–12% Bis-Tris pre-cast gel (Thermo Fisher) and transferred onto a PVDF membrane using an iBlot 2 gel transfer system (Thermo Fisher), following the manufactureŕs guidelines. Residual proteins remaining in the gel were visualized using SimpleBlue SafeStain (Thermo Fisher) and served as a loading control. The membrane was blocked in 5% milk in PBS with 0.1% Tween 20 (PBS-T) for 1 h at room temperature (RT) before being probed with either rat anti-HA-tag antibody (Roche, 11867423001) diluted 1:1000 in 5% milk in PBST, mouse anti-FLAG-M2-tag antibody (Sigma–Aldrich, F3165) diluted 1:2000 in 5% milk in PBST or mouse OXPHOS cocktail (abcam, ab110412) diluted 1:300 in 5% milk in PBST. After three washes with PBS-T for 10 min each at RT, the membranes were incubated with HRP-linked secondary antibodies, either anti-rat IgG (Cell Signaling, 7077 S) or anti-mouse IgG (Promega, W4021), diluted 1:5000 in 5% milk in PBST. Following another three washes as before, the membranes were digitally imaged using an Amersham Imager 680 blot and gel imager (GE Healthcare) upon exposure to Amersham ECLTM Western Blotting Detection Reagents (GE Healthcare).

### Results and discussion Adenine base editing *in vitro*

The originally reported adenine base editors (TALEDs) were shown to work within varying efficacies dependent on the architecture used [26]. The authors used either: (1) dimeric TALEDs (dTALED), which combines two sequence specific TALE arrays binding to opposite strands, with either the catalytically inactive (E1357A) DddAtox or TadA8e at their C-terminus, (2) split DddA TALED (sTALED) which uses the dimeric approach but with the DddAtox(E1357A) split at the G1397 position or (3) monomeric TALED (mTALED) which uses a single TALE-array attached to both the DddAtox(E1357A) and the TadA8e adenine deaminase (**Fig. 1a; Supplementary Fig. 1**). In order to test the efficiency of the TALED technology, we used validated TALE arrays for mtDNA-encoded mouse *Mt-Cytb*, *Mt-CoII* and *Mt-Atp6* from our previous work[13]. These TALEs are all known to bind the desired mtDNA position and induce high levels of cytosine base edits, and therefore acted as a good starting point to test the adenine base editing technology *in vitro*. Each of the TALE pair was cloned into plasmids containing the different DddAtox(E1357A)/TadA8e variations. This resulted in a total of 6 combinations which were transfected into WT NIH-3T3 mouse cells, sorted and replated to grow for 6 days. The levels of detectable editing from Sanger sequencing varied significantly (**Supplementary Figs. 2-4**), which was quantified using next generation sequencing (NGS) (**Fig. 1b**). For *Mt-Cytb*, low editing levels were observed in one of the dimeric DddAtox(E1357A) and TadA8e combinations, and also for the monomeric H strand binding construct (**Fig. 1b**). In the case of *Mt-CoII*, the dimeric DddAtox and TadA8e version showed editing alongside the split DddAtox(E1357A) dimeric combination (**Fig. 1b**). Finally, *Mt-Atp6* showed similar editing in both dimeric, split DddAtox(E1357A) and monomeric architectures (**Fig. 1b**). The highest editing efficiency we achieved after 7 days in vitro was 17%, comparable to previous reports for mouse NIH-3T3 cells, albeit lower than that observed in human cell A-to-G editing [26–28]. In the cases where editing was observed, it was present at multiple A:T sites within/across the editing window rather than being specific to predominantly one position as with the DdCBE technology[13]. The NGS analysis also allowed for mtDNA-wide off-target editing frequency to be calculated. The off-target editing was approximately 0.1% for all constructs which falls within the variation in unedited WT cells (**Fig. 1c**), suggesting generally lower activity of TALEDs, as compared to DdCBEs [13]. The off-targets for these constructs are therefore not concerning, although the on-target levels are relatively low and if they increased, the off-targets would likely follow.

**Fig. 1.**
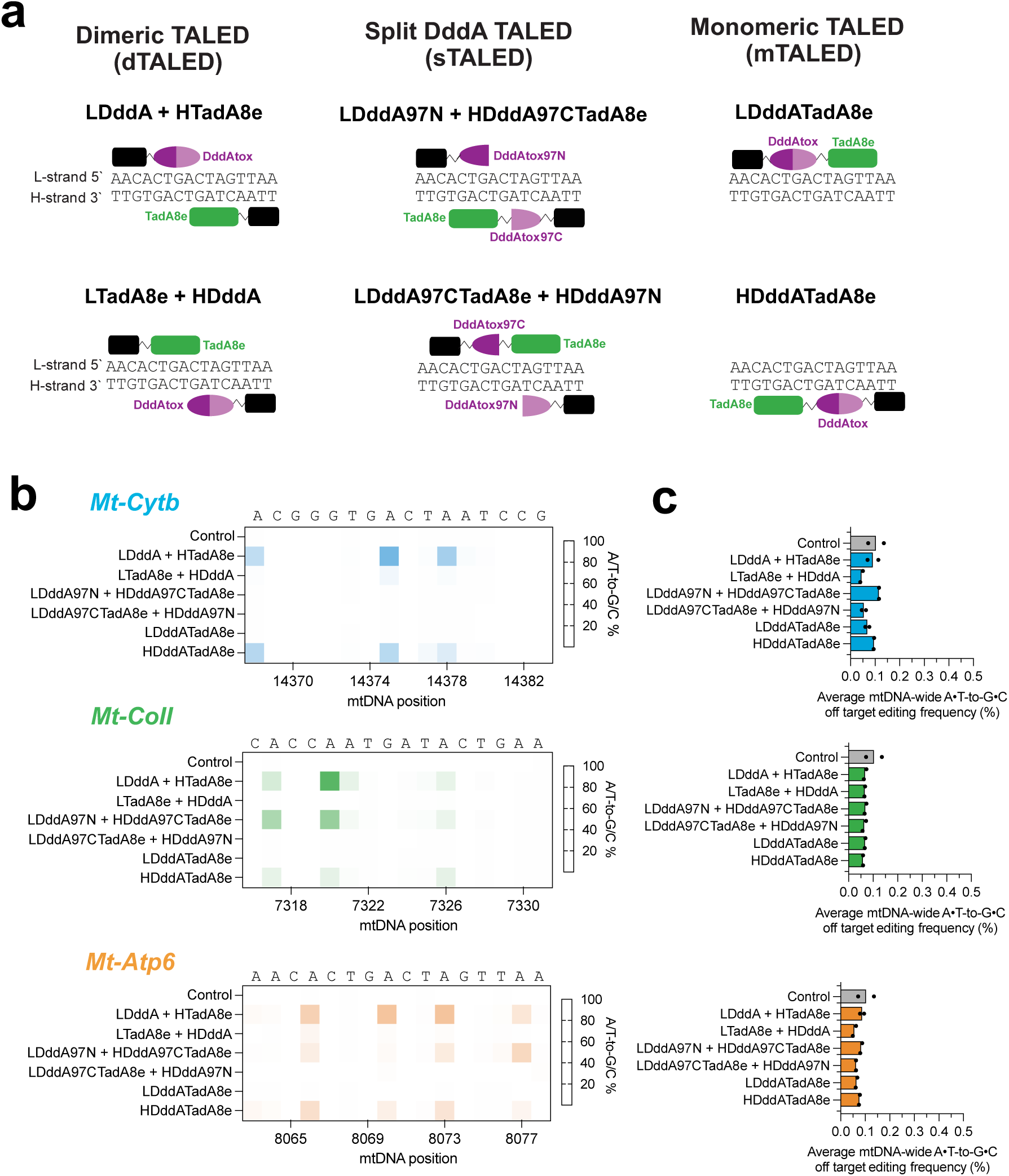
Mitochondrial adenine base editing by TALED in cultured mouse cells. **a** Schematic overview of the TALED adenine base editor architectures tested. The TadA8e enzyme is responsible for A:T to G:C editing, whilst DddAtox and Nt.BspD61(C) make the DNA accessible for enzyme activity. Black boxes indicate MTS and DNA binding elements. For full schematic see **Supplementary Fig. 1**. **b** On-target editing percentage values calculated by next generation sequencing of the six adenine base editor combinations targeted to the *Mt-Cytb*, *Mt-CoII* and *Mt-Atp6* genes using dTALED, sTALED and mTALED. Sequence and mouse mtDNA position is indicated. n = 2. **c** Average mtDNA-wide off-target editing frequency values calculated by next generation sequencing of the adenine base editor combinations targeted to the *Mt-Cytb*, *Mt-CoII* and *Mt-Atp6* genes from b. n = 2.

The observed strand bias (**Fig. 1**) could be attributed to TALED-mediated A-to-G editing being coupled to mitochondrial base-excision repair, in which DddAtox-dependent cytosine deamination promotes transient exposure of single-stranded DNA that can be accessed by TadA8e [29]. The DddAtox variant used here preferentially deaminates cytosines in a TC sequence context, and all three target regions contain a TC motif near the centre of the spacer on the H strand, making this strand particularly susceptible to DddAtox activity and thereby preferentially exposing the complementary L strand to TadA8e-mediated editing. Thus, the distribution and accessibility of DddAtox-susceptible cytosines [30], together with TALE orientation, are expected to influence which strand is edited. Consistent with this model, editing was predominantly observed on the L strand and was strongest when TadA8e was carried by the H-strand-binding monomer. We therefore favour an architecture- and sequence-dependent explanation rather than an intrinsic preference of TadA8e for either mtDNA strand.

We next decided to focus on *Mt-Atp6*, as we previously successfully edited this gene *in vitro* and *in vivo* using cytosine base editors[13]. We tested the strand selective mitochondrial base editors (mitoBEs), which use nickase enzymes to create the single stranded DNA required for TadA8e activity (**Fig. 2a, Supplementary Fig. 1**) [27]. We used the Nt.BspD61(C) nickase due to its lack of sequence specificity, unlike the MutH* nickase used in the previous study, which requires a 5’-GAT-3’ sequence not present in the *Mt-Atp6* target window. Following the NIH-3T3 cells transfection and selection (see above), the NGS analysis revealed up to 4% editing at multiple sites withing the targeting window between the TALE biding sites when using the mitoBEs (**Fig. 2b, Supplementary Fig. 5**), whilst mtDNA-wide off-targets were not significantly above control levels (**Fig. 2c**). Taken together, our study replicated the outcomes documented for both TALEDs and mitoBEs, with the efficiencies and editing patterns observed in mouse cells being largely consistent with those reported previously[26–28]. The lower activity of the adenine editors tested here compared with previously reported DdCBEs may in part reflect the distinct substrate requirements of their respective deaminases. DddA acts on double-stranded DNA, whereas TadA8e preferentially deaminates adenosines in single-stranded DNA. A-to-G editing therefore depends on architecture-specific generation or exposure of an accessible single-stranded DNA substrate. In mitoBEs this is promoted by strand nicking [27], while recent mechanistic work indicates that base-excision-repair-dependent DNA processing can enhance TALED activity [29]. This additional dependence on DNA processing likely contributed to the higher levels of editing produced by TALEDs when compared to mitoBEs (**Fig 2**).

**Fig. 2.**
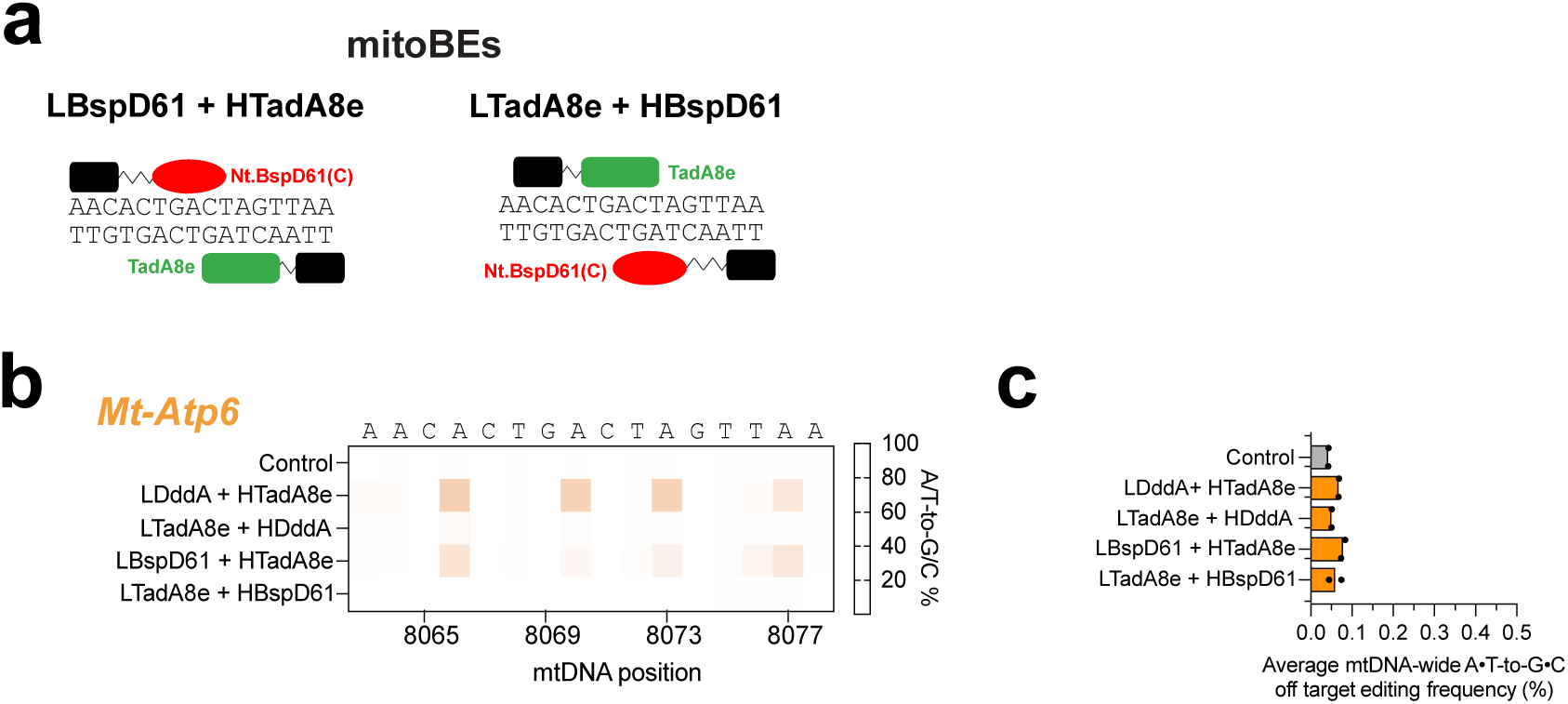
Mitochondrial adenine base editing by mitoBE in cultured mouse cells. **a** Schematic overview of the mitoBE adenine base editor architectures tested. The TadA8e enzyme is responsible for A:T to G:C editing, while Nt.BspD61(C) make the DNA accessible for editing activity. Black boxes indicate MTS and DNA binding elements. For full schematic see **Supplementary Fig. 1**. **b** On-target editing percentage values calculated by next generation sequencing of base editors targeted to *Mt-Atp6* gene using mitoBE, containing the Nt.BspD61(C) nickase. n = 2. **c** Average mtDNA-wide off-target editing frequency values from d. Multiple t test comparison performed.

### Adenine base editing *in vivo*

Next, we investigated whether mitochondrial adenine base editing could be achieved in somatic tissues following systemic AAV9 delivery to neonatal mice, using conditions similar to those employed previously for mitochondrial cytosine editors [20, 25]. Informed by our in vitro data, we selected the best-performing Mt-Atp6 constructs in the dimeric architecture, combining the H-strand-binding TadA8e construct with either DddAtox(E1357A) (the TALED architecture) or the Nt.BspD61(C) nickase (the mitoBE architecture) bound to the L strand. We cloned the respective sequences into AAV-compatible vectors and proceeded with encapsidation using our standard AAV production approach. However, titres for the H-strand TadA8e construct were approximately 1000-fold lower than for the L-strand DddAtox(E1357A) or Nt.BspD61(C) constructs and were insufficient for mouse injections; this outcome was reproduced at two independent vector-production facilities. During these experiments, a study reported extensive TadA8e-mediated RNA off-target editing associated with reduced cell viability. We therefore introduced the V28R substitution, reported to reduce RNA off-target activity. Consistent with the published findings, unmodified TadA8e caused substantial cell loss after transfection, whereas TadA8e(V28R) markedly improved cell survival (**Supplementary Fig. 6**) and enabled production of AAV at titres sufficient for in vivo experiments.

Neonatal mice were administered Mt-Atp6 H-strand TadA8e(V28R) AAV9 together with either DddAtox(E1357A) or Nt.BspD61(C) AAV9 by temporal-vein injection at 5 × 10^11 vg per monomer (**Fig. 3a**). At 4 weeks after administration, robust editor expression was detected in cardiac and skeletal muscle (**Supplementary Figs. 7** and **8**). Nevertheless, mtDNA editing remained low, reaching approximately 0.5% at the most efficiently edited positions in heart and remaining at or close to background in skeletal muscle and liver (**Fig. 3b**). To determine whether prolonged editor exposure could increase editing efficiency, a separate cohort was analysed 6 months after neonatal AAV administration. A-to-G editing increased modestly in animals receiving the TALED architecture, reaching approximately 1% at the most efficiently edited positions in heart and lower levels in skeletal muscle, whereas editing remained minimal in liver and in animals receiving the mitoBE architecture (**Fig. 3c**). Thus, the exposure period from 4 weeks to 6 months produced only a modest increase in editing, indicating that the low in vivo activity of these architectures cannot be explained simply by insufficient duration of editor expression. mtDNA-wide A:T-to-G:C editing remained at background levels for either 4-week or 6-month experiment (**Fig. 3d-e**), and no significant changes in mtDNA copy number or OXPHOS protein steady-state levels were detected upon 4-week treatment (**Fig. 3f** and **Supplementary Figs. 9** and **10**). Given the small group sizes, these *in vivo* data should be interpreted as molecular proof-of-concept rather than as powered estimates of tissue-specific editing efficiency.

**Fig. 3.**
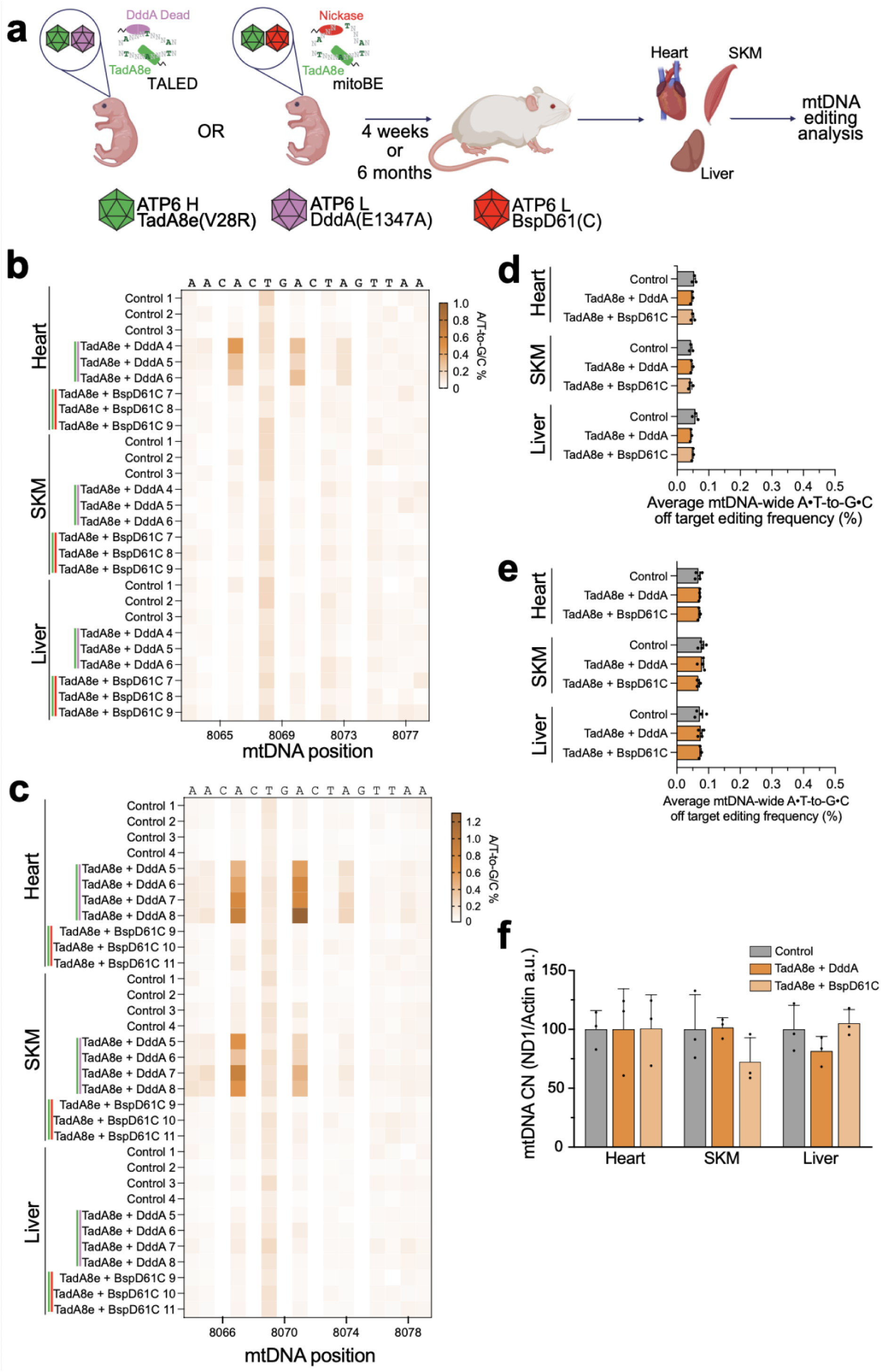
*In vivo* mouse mtDNA editing using adenine base editors. **a** Scheme of *in vivo* experiments with neonatal mice. TadA8e(V28R), DddAtox and BspD61(C) were encoded in separate AAV genomes, encapsidated in AAV9 then each pair (TadA8e(V28R) + DddAtox or TadA8e(V28R) + BspD61(C)) was simultaneously administered by temporal vein injection at 5 × 10^11^ vg/mouse of each monomer. Animals were sacrificed 4-weeks post-injection and their heart, skeletal muscle and liver tissues were examined for mtDNA editing. **b-c** On-target editing percentage values calculated by next generation sequencing of mouse tissues analysed following adenine base editor AAV delivery to the *Mt-Atp6* gene for 4 weeks (c) or 6 months (d). Sequence and mouse mtDNA position is indicated. **d-e** Average mtDNA-wide off-target editing frequency values calculated by next generation sequencing from mice in treated for 4 weeks (d) or 6 months (e). Bars represent the mean and error bars represent ± SEM (n = 3). **f** Real-Time qPCR relative quantification of mtDNA copy number (CN) in 4-week-old mouse tissues following adenine base editor AAV delivery. Bars represent the mean and error bars represent ± SEM (n = 3). Multiple t test comparison performed.

### Conclusions and future directions

In this study, we tested the effect of adenine base editors first in cells using our TALEs targeted to *Mt-CytB, Mt-CoII* and *Mt-Atp6* and confirmed editing within the target window. We then tested the effect of *Mt-Atp6* adenine base editors in mice using AAV delivery to neonate mice and observed negligible editing in mouse hearts, with no editing in other tissues. In comparable experimental conditions, TALE-based DdCBE and ZF-DdCBEs installed up to 30% and 50% of C:G to T:A edits, respectively [20, 25]. Considering these limitations, the tested adenine base editor architectures need further optimization to become a valuable tool for generating or correcting pathogenic *in vivo* point pathological variants associated with mitochondrial diseases. The present study, therefore, establishes feasibility but also defines clear requirements for further development. Developments reported since this study was initiated demonstrate that these limitations are tractable. More specifically, optimised mitoBEv2 architectures have achieved high-efficiency and precise mitochondrial A-to-G editing in mouse disease models [31]; enhanced TALED architectures modulate mtBER activity to improve editing activity and precision [29]; and evolved TadA8e [29, 32] and DddAtox [29] variants substantially increased activity across diverse sequence contexts while reducing DNA and RNA off-target effects. Future mitochondrial adenine editors for in vivo use should combine high on-target activity with narrow or ideally single-nucleotide editing windows, minimal mtDNA-wide, nuclear-DNA and RNA off-target activity, low cellular toxicity, compatibility with practical delivery systems, and efficient activity in mature disease-relevant tissues, including central nervous system.

## Data availability

The data supporting the findings of this study are available within the paper and its supplementary information files, apart from proprietary scripts which are available upon request by contacting the authors. The NGS files generated in this study have been deposited in GEO: GSE276406: Source data are provided with this paper. The materials used in this study are available upon request from M.M.

## Supplementary data

Supplementary Data are attached to this manuscript.

## Acknowledgments

We would like to acknowledge the members of the Mitochondrial Genetics Group (MRC-MBU, University of Cambridge) for useful discussion during the course of this research. We would also like to thank the CIMR Flow facility for their support in this work.

## Authors’ contributions

C.D.M., P.S.-P. and M.M. planned and designed experiments; C.D.M. performed most of the experiments; L.V.H. analysed the NGS experiments; L.L.G. and I.D. were involved in the NGS experiments; P.S.-P. performed the AAV in vivo experiments; K.T. managed the animal work; J.D.B-P help interpreting the data; C.D.M., P.S.-P. and M.M. drafted the manuscript. P.S.-P. and M.M. supervised the study and acquired funds. All authors revised the manuscript.

## Supplementary material

**Supplementary Figure 1:**
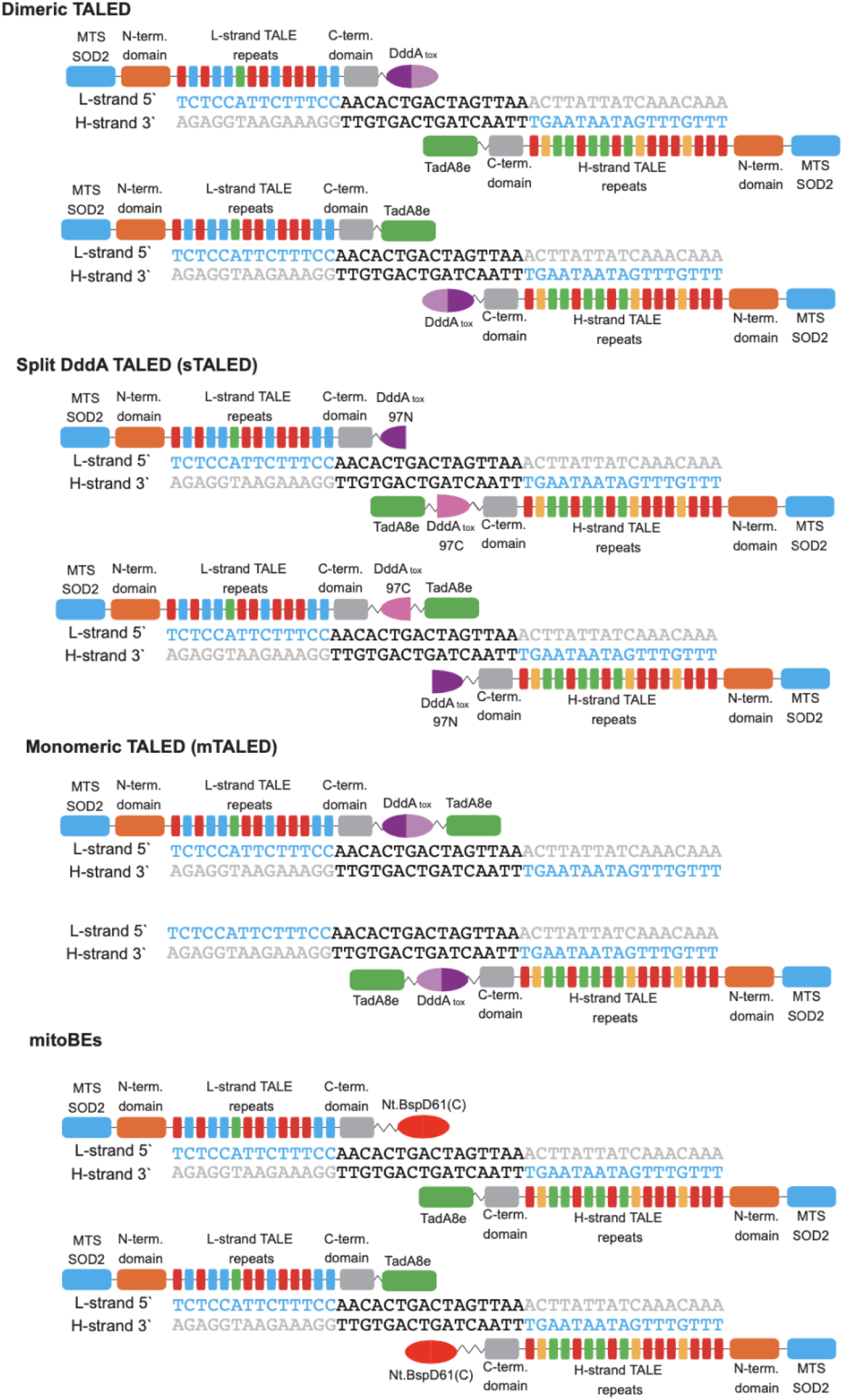
Strategies for adenine base editing. Schematics showing the four main methods used for adenine base editing in mitochondria. **Dimeric TALED (dTALED)** uses a TALE-binding mtDNA sequence specific protein attached to the catalytically dead DddAtox (for DNA unwinding) on one monomer, and the TadA8e adenine deaminase to the monomer binding the opposite mtDNA strand. **Split DddA TALED (sTALED)** which uses the G1397 split of the DddAtox attached to two TALED monomers, alongside the TadA8e. **Monomeric TALED (mTALED)** which contains only one sequence specific TALE protein attached to both the DddAtox and the TadA8e. Dimeric **mitoBEs** use nickase enzyme BspD61(C) to allow for single stranded activity of TadA8e.

**Supplementary Figure 2:**
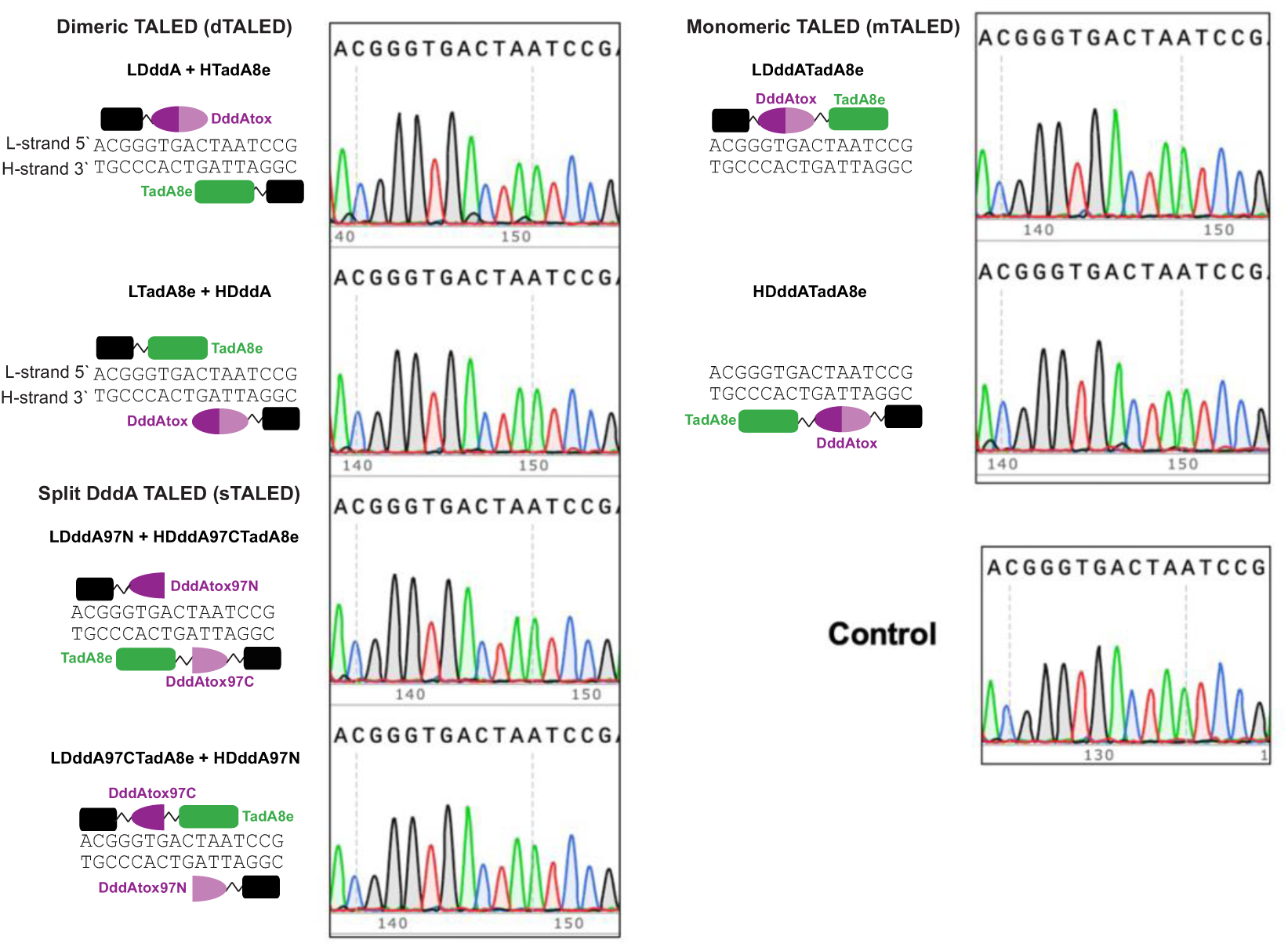
Adenine base editing of *Mt-Cytb*. Sanger sequencing traces of the six adenine base editor combinations targeted to the *Mt-Cytb* gene. Only the on-target editing window is shown.

**Supplementary Figure 3:**
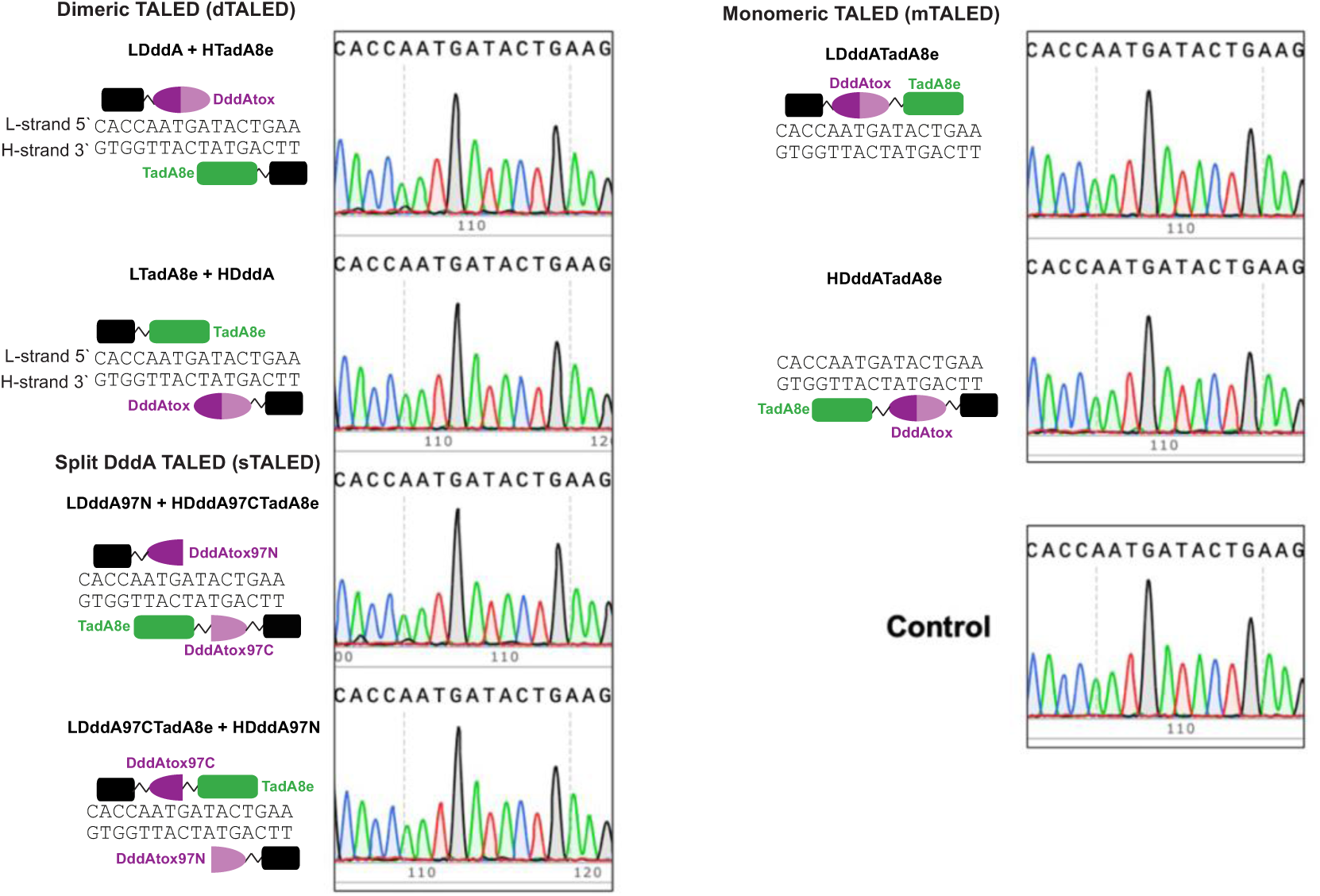
Adenine base editing of *Mt-CoII*. Sanger sequencing traces of the six adenine base editor combinations targeted to the *Mt-CoII* gene. Only the on-target editing window is shown.

**Supplementary Figure 4:**
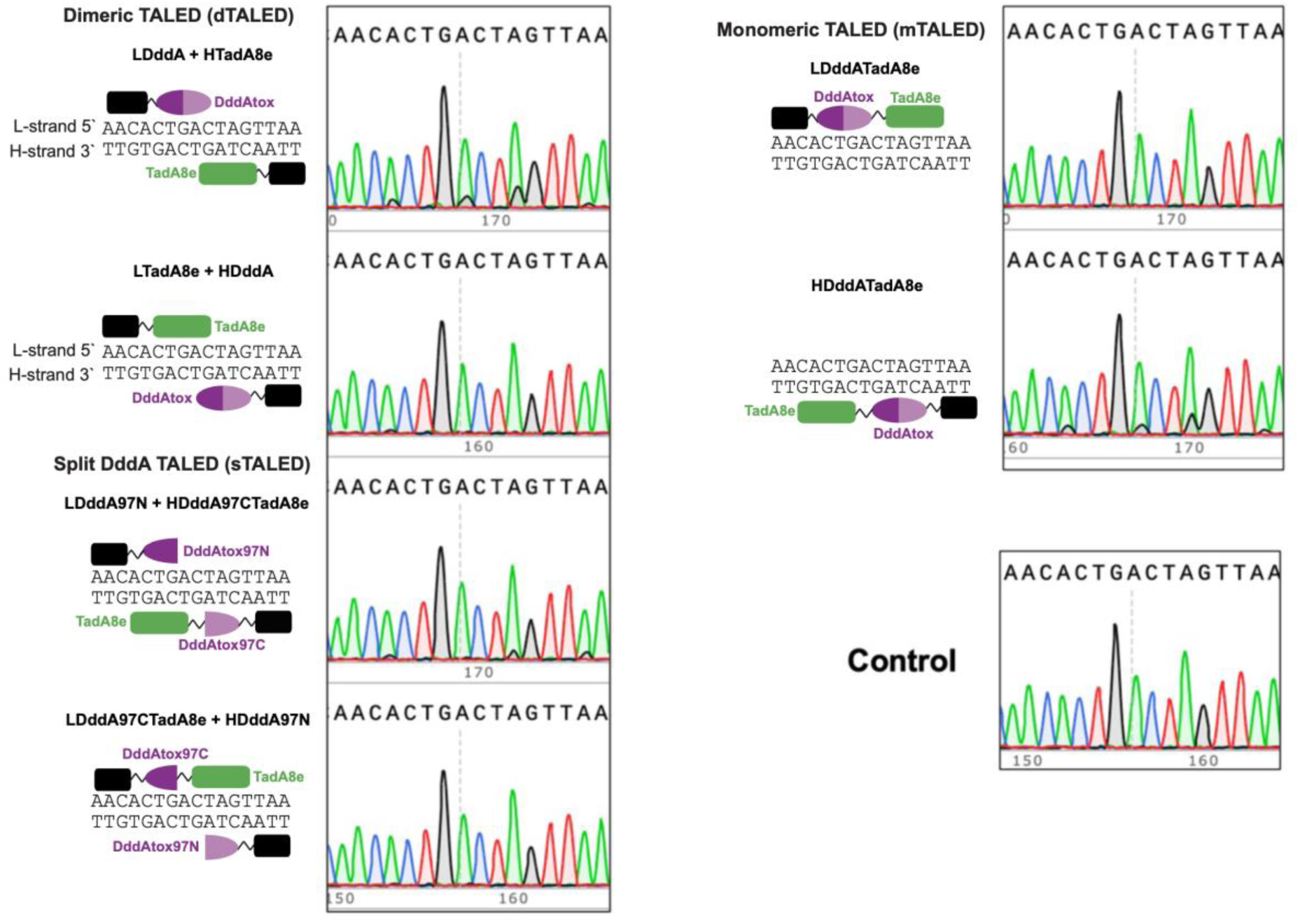
Adenine base editing of *Mt-Atp6*. Sanger sequencing traces of the six adenine base editor combinations targeted to the *Mt-Atp6* gene. Only the on-target editing window is shown.

**Supplementary Figure 5:**
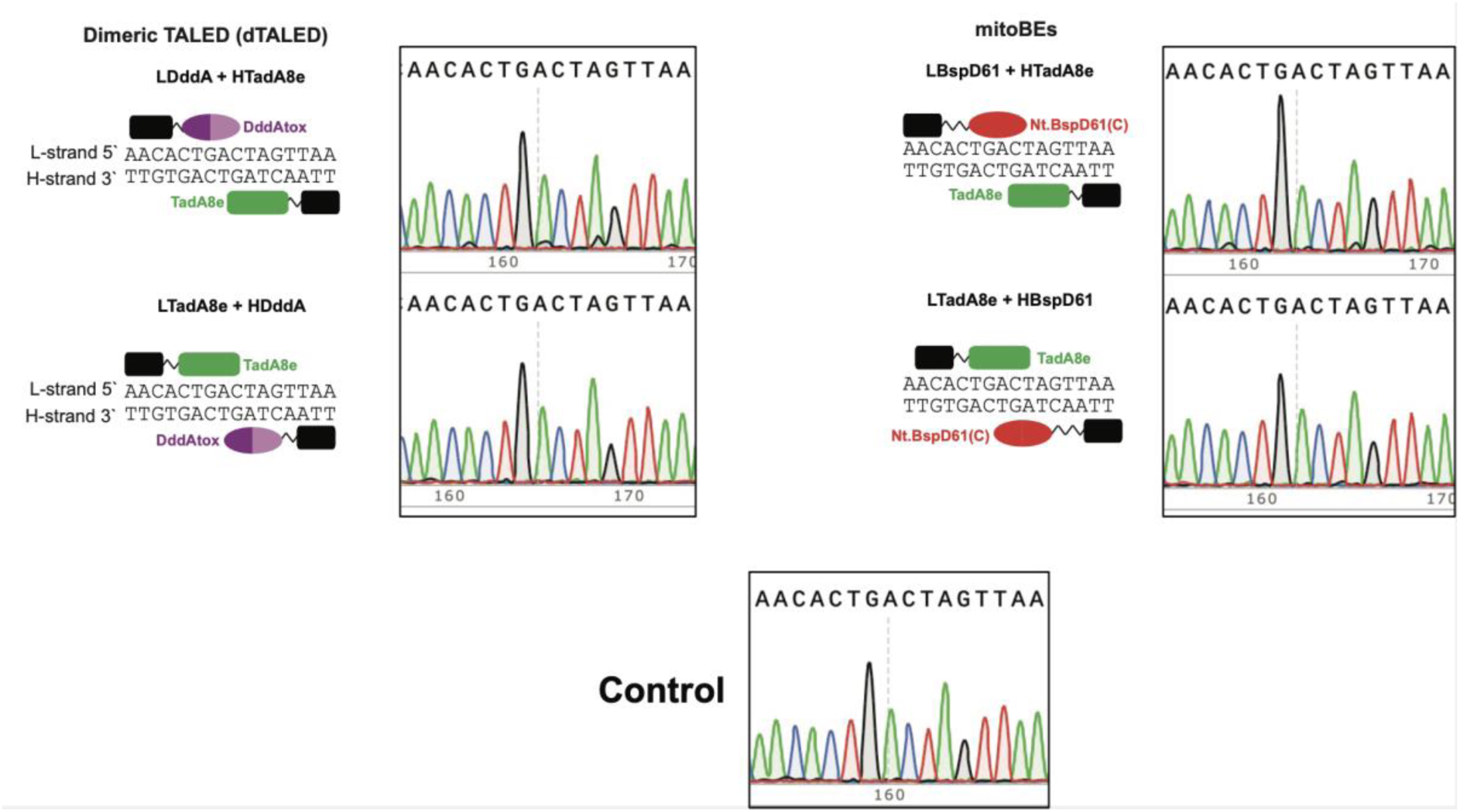
Adenine base editing of *Mt-Atp6* by mitoBEs. Sanger sequencing traces for the two mitoBE combinations targeting to the *Mt-Atp6* gene, compared to dTALED editing directed to the same gene. Only the on-target editing window is shown.

**Supplementary Figure 6:**
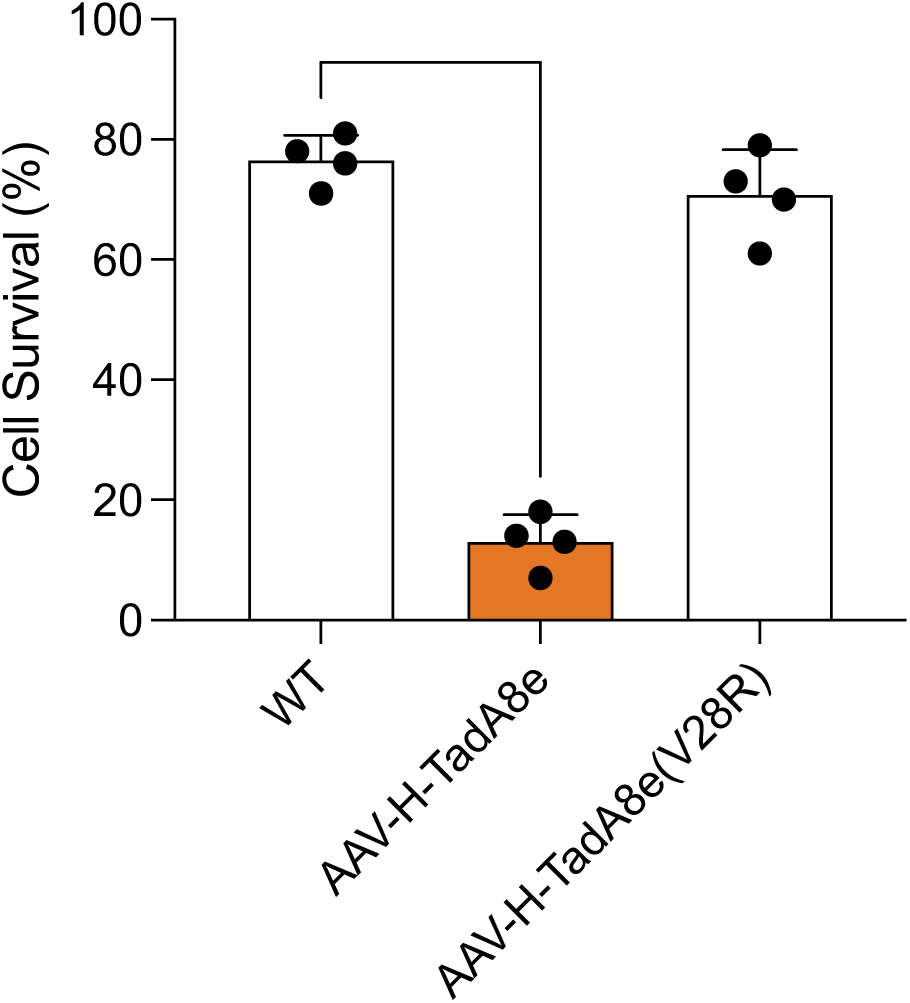
Cell survival following transfection with AAV plasmids encoding TadA8e variants. Cell survival percentage calculated using Countess Automated Cell Counter following 4 days post transfection of AAV plasmids spiked with empty GFP plasmid and sorting for enrichment. n = 4 biological replicates. Error bars indicate ±SEM Statistical significance was calculated from a one-tailed Student’s t test **** P-value < 0.0001.

**Supplementary Figure 7:**
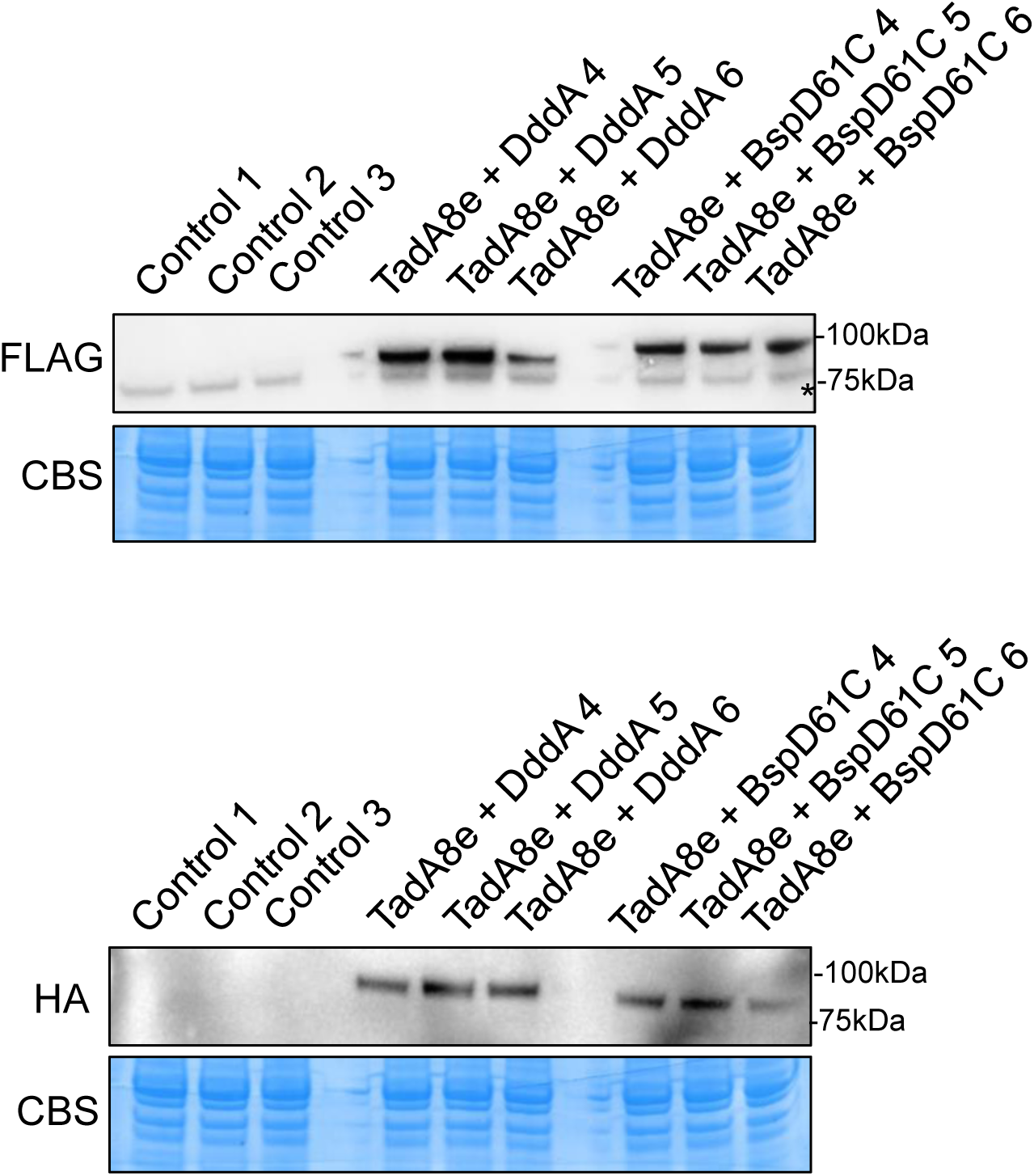
Delivery of adenine base editor AAVs to mouse skeletal muscle. Western blot analysis of the levels of FLAG-tagged AAV DddA or BspD61C (L) and HA-tagged AAV TadA8e (H) in skeletal muscle (quadriceps) of mice at 4 weeks post-injection. Coomassie Brilliant Blue (CBB) was used as loading control. * indicates a non-specific band detected by the anti-FLAG antibody.

**Supplementary Figure 8:**
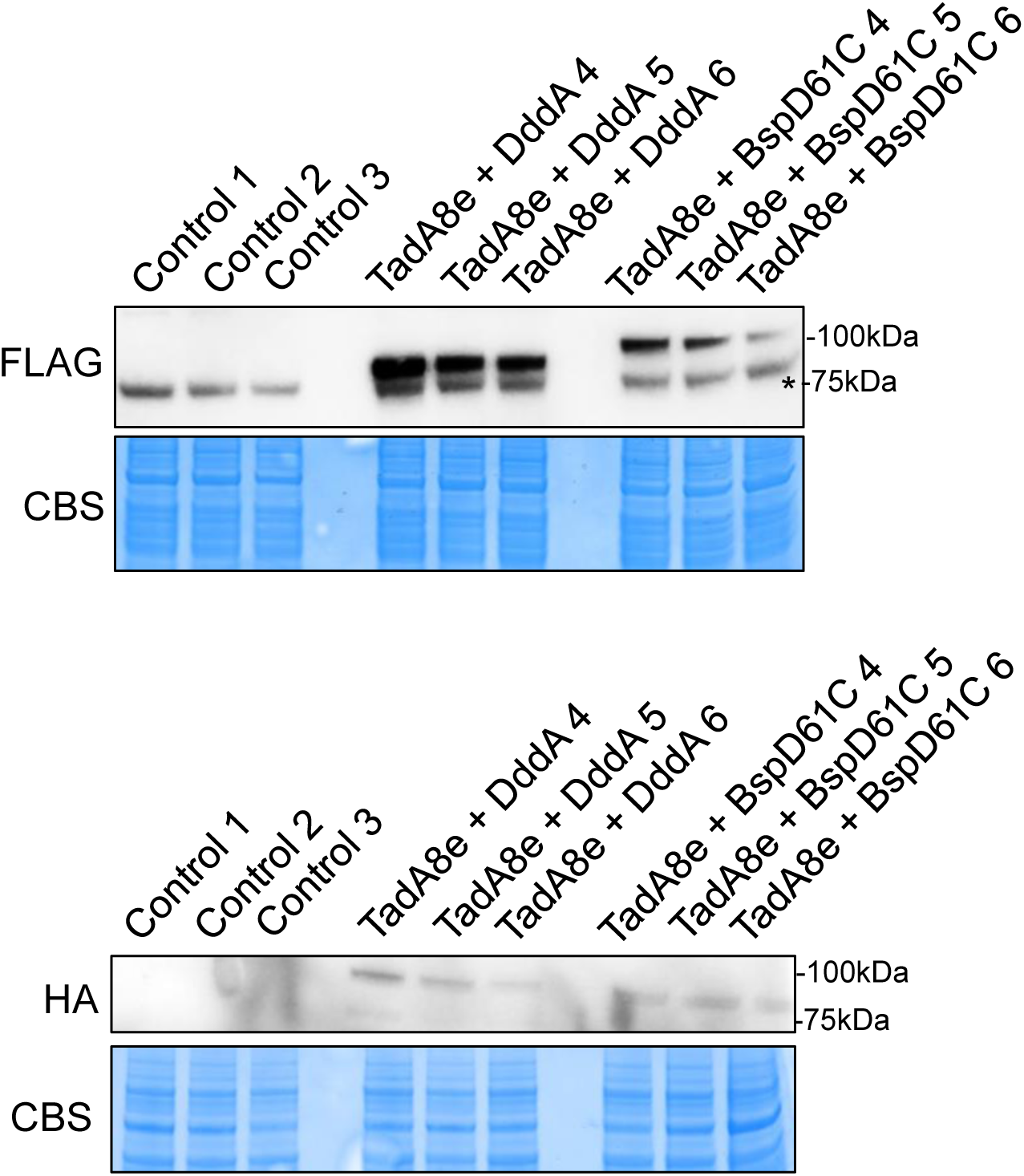
Delivery of adenine base editor AAVs to mouse heart. Western blot analysis of the levels of FLAG-tagged AAV DddA or BspD61C (L) and HA-tagged AAV TadA8e (H) in heart of mice at 4 weeks post-injection. Coomassie Brilliant Blue (CBB) was used as loading control. * indicates a non-specific band detected by the anti-FLAG antibody.

**Supplementary Figure 9:**
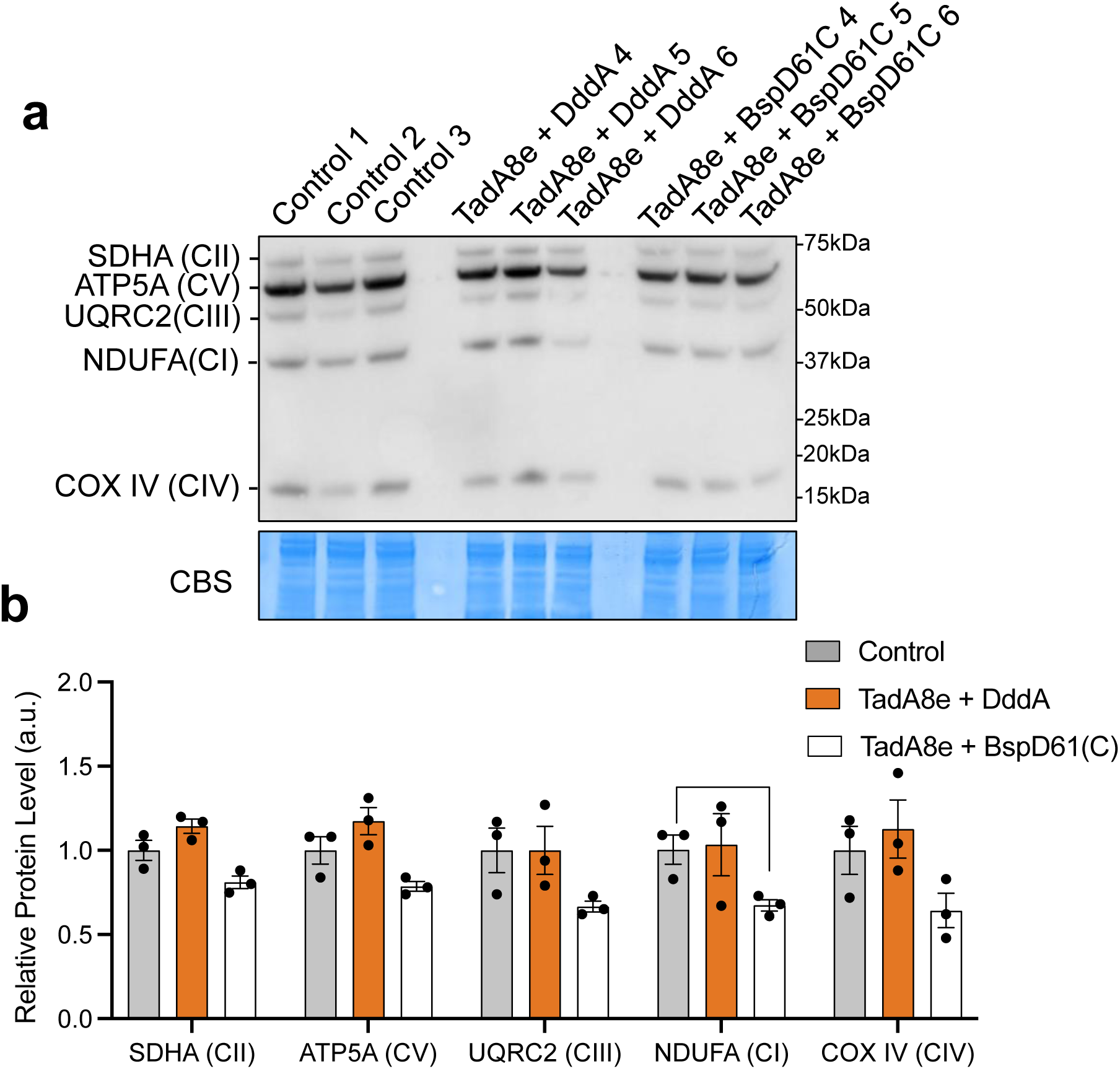
Assessment of OXPHOS complex levels in skeletal muscle following adenine base editor AAV delivery. **a** Western blot analysis of steady-state levels of components of OXPHOS in skeletal muscle of 4-week old control mice, mice injected with AAVs for TadA8e + DddA or TadA8e + BspD61(C). Coomassie Brilliant Blue (CBB) was used as loading control. **b** The western blots from a quantified using ImageJ. Error bars indicate ±SEM. Multiple t test comparison performed. *p<0.05.

**Supplementary Figure 10:**
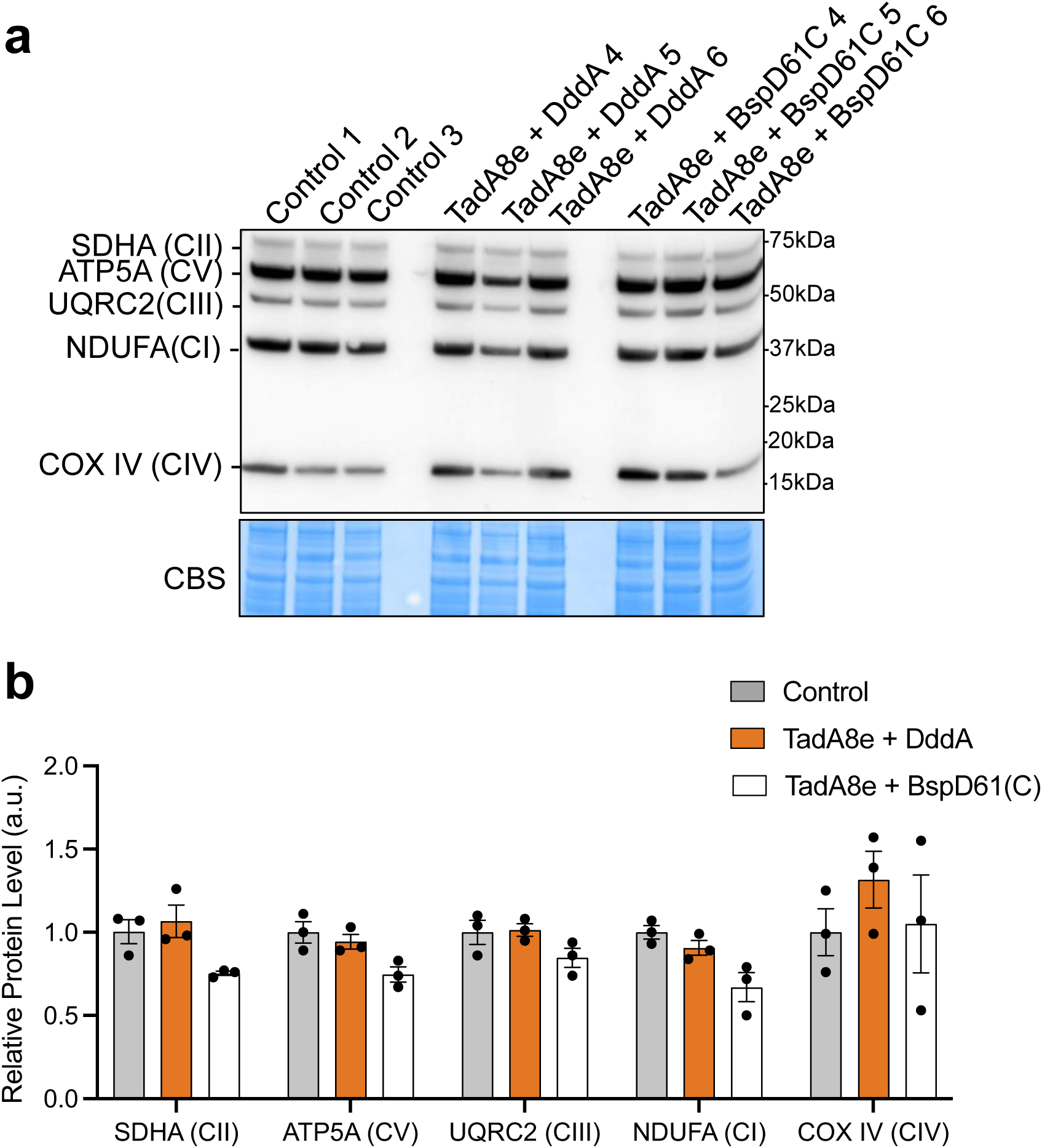
Assessment of OXPHOS complex levels in heart following adenine base editor AAV delivery. **A** Western blot analysis of steady-state levels of components of OXPHOS in heart of 4-week old control mice, mice injected with AAVs for TadA8e + DddA or TadA8e + BspD61(C). Coomassie Brilliant Blue (CBB) was used as loading control. **b** The western blots from a quantified using ImageJ. Error bars indicate ± SEM. Multiple t test comparison performed.

